# Cellular type is a major determinant of R-loop genomic distribution

**DOI:** 10.1101/2025.04.17.649299

**Authors:** Katerina Oleynikova, Nadezhda Zhigalova, Andrew P Hutchins, Alexey Ruzov

## Abstract

R-loops containing a DNA:RNA hybrid and unpaired single-stranded DNA are important for normal cell physiology and pathogenesis of numerous diseases. Although several different approaches for R-loop mapping have been developed, these techniques can produce conflicting results. In order to assess their robustness, a study by Chedin et al compared a number of R-loop datasets obtained using different methodologies from normal and cancer cell lines. That study assumed a high degree of similarity between R-loop genomic distributions across different cell types. Here, comparing DRIP datasets produced using the same protocol in different cell lines; we show that only 26 percent of R-loop peaks are common between a chronic myeloid leukemia-derived HAP1 and human pluripotent stem cells, whereas HAP1-derived knockout cell lines share substantial fractions of R-loop peaks with their parental line. We conclude that cellular type represents a major determinant of R-loop genomic distribution and, therefore, only a systematic comparison of a large array of various cell/tissue-types-derived R-loop datasets may address the inconsistencies between different R-loop mapping techniques.

---

R-loops are three-stranded nucleic acid structures that contain a DNA:RNA hybrid formed by annealing of the nascent RNA molecule to the template DNA together with the displaced single-stranded DNA (García-Muse and Aguilera, 2019). R-loops are abundant in the genome and, depending on the biological context, may be involved in the regulation of a wide range of crucial biological processes such as transcription termination, cohesin-mediated DNA looping, DNA repair, telomere homeostasis and Immunoglobulin class switching recombination (Aguilera and Aguilera, 2025). At the same time, non-scheduled or pathological R-loops may interfere with transcription and replication leading to accumulation of double-stranded DNA breaks and, thus, representing a major source of genetic stress and instability in mammalian cells (García-Muse and Aguilera, 2019). Given the association of genome instability with cancer and certain genetic diseases such as neurodegenerative conditions and cancer-prone syndromes, understanding the regulation and genomic distribution of R-loops in different model systems has attracted considerable research attention, and a large array of experimental techniques that map R-loops in the genome have been developed (Yadav et al, 2024). Some of these methods are based on the use of an antibody that specifically recognizes RNA:DNA hybrids (S9.6 antibody) (Boguslawski et al, 1986) and include DNA:RNA Immunoprecipitation (DRIP) together with its numerous variants (e.g. DRIPc-seq; single-stranded DNA adaptor ligation DRIP, ssDRIP-seq etc) and R-Loop Cleavage Under Targets and Tagmentation (R-Loop CUT&Tag) (Ginno et al, 2012; García-Rubio et al, 2022; Sanz and Chedin, 2019; Wang et al, 2022; Crossley and Cimprich, 2022; Vachez et al, 2022; Xu et al, 2022). Other techniques rely either on bisulfite-based mapping of single stranded DNA R-loop components (Malig et al, 2020) or on the use of RNase H1, the enzyme responsible for recognition and eventual resolution of RNA:DNA hybrids (Chen et al, 2017; Cerritelli et al, 2022; Wulfridge et al, 2022; Zhang et al, 2022). Importantly, despite the wide selection of the mapping techniques, R-loop datasets produced using different (and even sometimes analogous) methodologies on various cell lines in different labs may often look rather dissimilar (Chedin et al, 2021).

Aimed at addressing these inconsistencies, a study by Chedin et al compares a number of R-loop mapping datasets obtained using different methods on various cell lines in order to assess the commonality of existing datasets (Chedin et al, 2021). It is worth noting that this paper contains a number of interesting insights into the nature of discrepancies between the results acquired using antibody-based and RNase H-based R-loop mapping techniques. However, it does not fully consider the cell types of the datasets in question. The implicit assumption is that there is a core similarity between R-loops at housekeeping genes between different cell types. However, it is not clear that this is a valid assumption, and it is possible, even likely, that R-loops are highly divergent between different cell types. Thus, the authors do not evaluate or discuss the potential degree of dis(similarity) between R-loop genomic distributions in biologically distinct cellular systems such as human pluripotent stem cells (hPSCs) and cancer-derived U2OS or HeLa cells and use the comparison of the signal coverage profiles of the corresponding DRIP experiments from different cellular types for the identification of the datasets they consider discordant. Hence, we believe that such an approach implicitly assuming that R-loop genomic distributions in cell lines of different origin are highly similar is not supported by experimental results and may lead to potential misrepresentation of experimental data.

To assess the level of similarity between R-loop genomic distributions in the cells of different origin, we compared the DRIP datasets produced using the same previously published protocol (Abakir et al, 2020) on three different cellular models: hPSCs, HAP1 “wild type” (HAP1 WT) cells derived from a chronic myeloid leukemia cell line KBM-7 (Carette et al, 2009) and two HAP1 isogenic knockout cell lines, with *YTHDF2* or *HNRNPA2B1* genes knocked out via genetic editing (HAP1 YTHDF2 KO and HAP1 2B1 KO cell lines, respectively). We performed DRIP-seq to determine the genomic regions enriched with R-loops in these cell lines, and after mapping reads to the human genome, we called peaks (Zhang et al, 2008), followed by identification of confident peaks for each sample by comparing the corresponding replicates (Supplementary Tables S1-S2, the code used for the analysis is available on GitHub: https://github.com/katerinaoleynikova/human_samples_paper_25). Our analysis revealed that although hPSCs and HAP1 WT DRIP datasets produced similar numbers of confident peaks (4482 and 4458 peaks correspondingly), only 1187 peaks (about 26%) were common between these two cell lines (Fig. 1A). Interestingly, whereas the knockouts of both *YTHDF2* and *HNRNPA2B1* resulted in a significant drop in overall peak numbers (1496 and 950 peaks correspondingly) compared with HAP1 WT cells, HAP1 YTHDF2 KO and HAP1 2B1 KO R-loop datasets were highly similar to each other with the overwhelming majority of HAP1 2B1 KO peaks (873 out of total 950) overlapping with those identified in HAP1 YTHDF2 KO cells (Fig. 1B). Moreover, 71% (681) of peaks identified in HAP1 2B1 KO cells and 55% (864) of HAP1 YTHDF2 KO peaks were identical to the R-loop peaks found in their parental HAP1 WT cell line (Fig. 1B). Furthermore, the genomic region centered around housekeeping gene *RPL13A* used by Chedin et al for their datasets comparison and designated as a “gold standard region”, contained only HAP1 WT peaks but not R-loop peaks from hPSCs or other two HAP1 knockout datasets tested in our study (Fig. 1C). Indeed, we identified other loci where R-loop peaks were present in all 4 of the tested cell lines (Fig. 1D left panel). However, in our opinion, the designation of gold standard regions requires generating datasets from other cell types as R-loops appear highly cell type-specific.

**Fig. 1.**
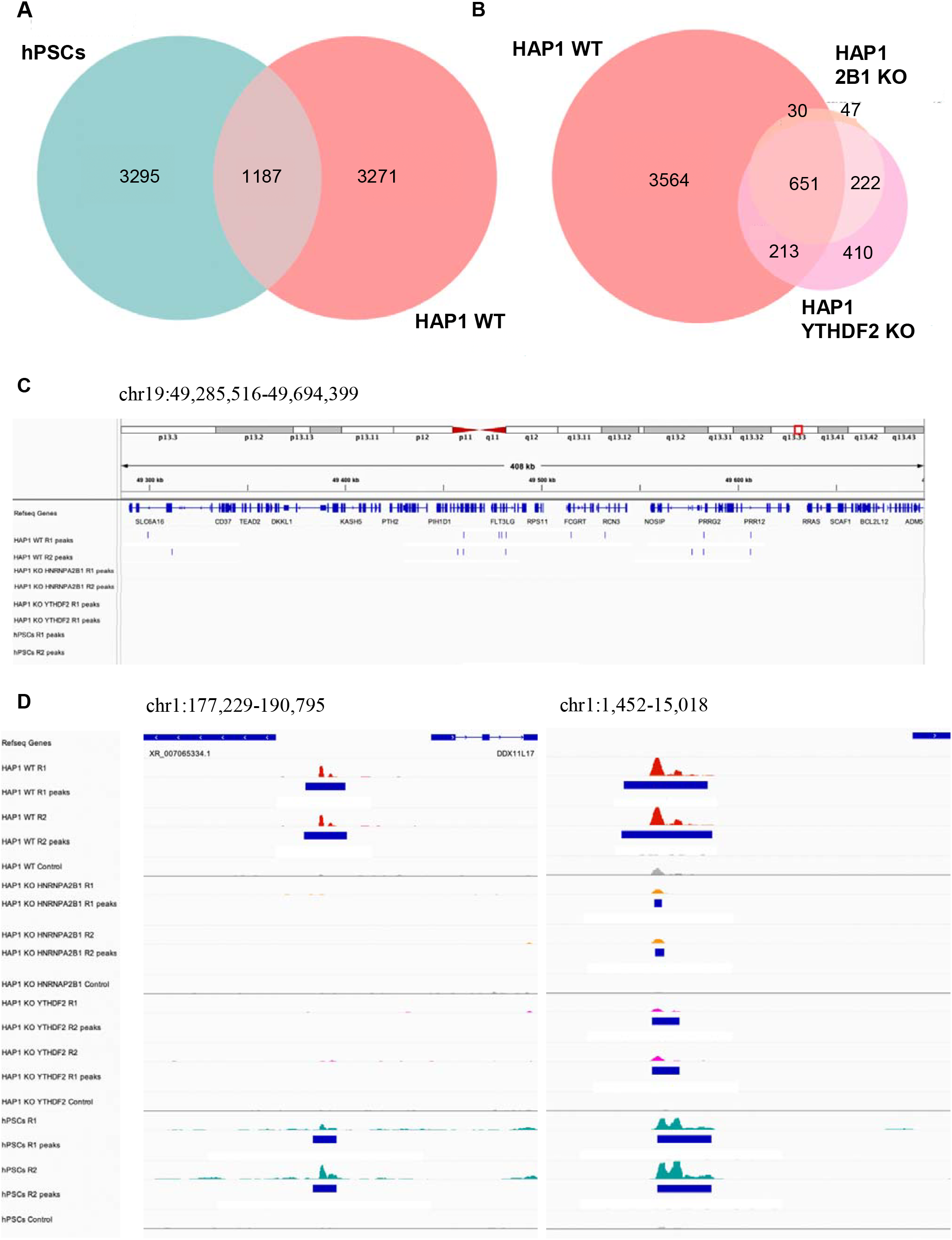
Cellular type is a major determinant of R-loop genomic distribution in human cells. **(A, B)** Venn diagrams showing the overlaps between the R-loop peak datasets obtained from DRIP experiments performed using our previously published protocol (Abakir et al, 2020) on hPSCs and wild type HAP1cells (HAP1 WT) (**A**), as well as wild type HAP1cells and HAP1 cells with genetic knockouts of YTHDF2 (HAP1 YTHDF2 KO) and HNRNPA2B1 (HAP1 2B1 KO) genes (**B**). The numbers of R-loop peaks in each category are indicated on the diagrams. (**C**) Genome browser view of the distribution of R-loop peaks (shown as blue vertical dashes) in the datasets generated in our DRIP experiments on the indicated cell lines (HAP1 wild type, HAP1 YTHDF2 KO, HAP1 HNRNPA2B1 KO and hPSCs) over the region centered around housekeeping gene RPL13A used by Chedin et al for their comparison and designated as “the standard “gold standard” in that study. Notably, among our tested datasets, this region contains only HAP1 wild type peaks and does not enclose any hPSCs R-loop peaks. (**D**) Genome browser views of signal profiles of our R-loop DRIP datasets alongside the control input samples at two representative genomic loci. The locations of the R-loop peaks are shown with blue rectangles. Whereas R-loop peaks are present only in HAP1 wild type and hPSCs cells in the locus showing in left panel, the right panel contains the site with R-loop peaks detectable in all the tested cell lines. Genome coordinates are shown for each region in (**C, D**).

Thus, our analysis shows that R-loop datasets produced using the same methodology substantially differ between cell lines of different origin with only around a quarter of R-loop peaks in common between hPSCs and a chronic myeloid leukemia-derived HAP1 cell line. At the same time the R-loop distributions in HAP1 knockout and wild type cells seem more similar to each other compared with that of hPSCs. Given that factors recently implicated in R-loop regulation include chromatin structure (Bayona-Feliu et al, 2021), nucleosome positioning (Abakir and Ruzov, 2021; Werner et al, 2025) and RNA modifications (Abakir et al, 2020; Abakir and Ruzov, 2024) that all vary widely between different cell types, these results do not seem particularly surprising for us. In summary, we conclude that R-loops are cell type-specific and, therefore, only a systematic comparison of a large array of various cell/tissue-types-derived R-loop datasets may address the inconsistencies between currently available R-loop mapping techniques and experimental results.

## Author contributions

A.R. conceived the paper, drafted and edited the manuscript and the figure. K.O. performed bioinformatics analyses. K.O. and A.P.H contributed to writing of the text, editing the manuscript and the figure preparation. N.Z. performed cell culture and molecular biology experiments. All authors read and approved the final manuscript.

## Acknowledgements

A.R.’s laboratory is funded by the Russian Science Foundation (grant no. 22-65-00022).

## Data availability

The hPSCs deep sequencing data can be found in the NCBI Sequence Read Archive (SRA) with the Bioproject ID: PRJNA474076 (https://www.ebi.ac.uk/ena/browser/view/PRJNA474076?show=reads). The HAP1-related deep sequencing datasets have been deposited in the NCBI SRA with the Bioproject ID: PRJNA1250978 (https://www.ncbi.nlm.nih.gov/bioproject/PRJNA1250978). The samples are labelled as: WT – HAP1 WT; KOYF2 - HAP1 YTHDF2 KO; KO2B1 – HAP1 HNRNPA2B1 KO. The in-house scripts used for the analysis have been deposited to GitHub online repository (https://github.com/katerinaoleynikova/human_samples_paper_25).

## Supplementary Material

Supplementary information contains Supplementary Tables S1-2 and Supplementary Materials and Methods.

## References

Abakir, A., Giles, T. C., Cristini, A., Foster, J. M., Dai, N., Starczak, M., … & Ruzov, A. (2020). N6-methyladenosine regulates the stability of RNA: DNA hybrids in human cells. Nature genetics, 52(1), 48–55.

Abakir, A., & Ruzov, A. (2021). SWI/SNF complexes as determinants of R-loop metabolism. Nature Genetics, 53(7), 940–941.

Abakir, A., & Ruzov, A. (2024). A model for a dual function of N6-methyladenosine in R-loop regulation. Nature Genetics, 56(10), 1995–1998.

Aguilera, P., & Aguilera, A. (2025). R-loop homeostasis in genome dynamics, gene expression and development. Current Opinion in Genetics & Development, 92, 102325.

Andrews, S., Krueger, F., Segonds-Pichon, A., Biggins, L., Krueger, C., & Wingett, S. (2010). FastQC. A quality control tool for high throughput sequence data, 370.

Bayona-Feliu, A., Barroso, S., Muñoz, S., & Aguilera, A. (2021). The SWI/SNF chromatin remodeling complex helps resolve R-loop-mediated transcription–replication conflicts. Nature genetics, 53(7), 1050–1063.

Carette, J. E., Guimaraes, C. P., Varadarajan, M., Park, A. S., Wuethrich, I., Godarova, A., … & Brummelkamp, T. R. (2009). Haploid genetic screens in human cells identify host factors used by pathogens. Science, 326(5957), 1231–1235.

Cerritelli, S. M., Sakhuja, K., & Crouch, R. J. (2022). RNase H1, the gold standard for R-loop detection. R-Loops: Methods and Protocols, 91–114.

Chédin, F., Hartono, S. R., Sanz, L. A., & Vanoosthuyse, V. (2021). Best practices for the visualization, mapping, and manipulation of R-loops. The EMBO Journal, 40(4), e106394.

Chen, L., Chen, J. Y., Zhang, X., Gu, Y., Xiao, R., Shao, C., … & Fu, X. D. (2017). R-ChIP using inactive RNase H reveals dynamic coupling of R-loops with transcriptional pausing at gene promoters. Molecular cell, 68(4), 745–757.

Crossley, M. P., & Cimprich, K. A. (2022). Quantitative DNA–RNA immunoprecipitation sequencing with spike-ins. In R-Loops: Methods and Protocols (pp. 381–410). New York, NY: Springer US.

García-Muse, T., & Aguilera, A. (2019). R loops: from physiological to pathological roles. Cell, 179(3), 604–618.

García-Rubio, M., Soler-Oliva, M. E., & Aguilera, A. (2022). Genome-Wide Analysis of DNA– RNA Hybrids in Yeast by DRIPc-Seq and DRIP-Seq. In R-Loops: Methods and Protocols (pp. 429–443). New York, NY: Springer US.

Ginno, P. A., Lott, P. L., Christensen, H. C., Korf, I., & Chédin, F. (2012). R-loop formation is a distinctive characteristic of unmethylated human CpG island promoters. Molecular cell, 45(6), 814–825.

Krueger, F. (2015). Trim Galore!: A wrapper around Cutadapt and FastQC to consistently apply adapter and quality trimming to FastQ files, with extra functionality for RRBS data. Babraham Institute.

Langmead, B., & Salzberg, S. L. (2012). Fast gapped-read alignment with Bowtie 2. Nature methods, 9(4), 357–359.

Li, H., Handsaker, B., Wysoker, A., Fennell, T., Ruan, J., Homer, N., … & 1000 Genome Project Data Processing Subgroup. (2009). The sequence alignment/map format and SAMtools. bioinformatics, 25(16), 2078–2079.

Malig, M., Hartono, S. R., Giafaglione, J. M., Sanz, L. A., & Chedin, F. (2020). Ultra-deep coverage single-molecule R-loop footprinting reveals principles of R-loop formation. Journal of Molecular Biology, 432(7), 2271–2288.

Quinlan, A. R., & Hall, I. M. (2010). BEDTools: a flexible suite of utilities for comparing genomic features. Bioinformatics, 26(6), 841–842.

Sanz, L. A., & Chédin, F. (2019). High-resolution, strand-specific R-loop mapping via S9. 6-based DNA–RNA immunoprecipitation and high-throughput sequencing. Nature protocols, 14(6), 1734–1755.

Thorvaldsdóttir, H., Robinson, J. T., & Mesirov, J. P. (2013). Integrative Genomics Viewer (IGV): high-performance genomics data visualization and exploration. Briefings in bioinformatics, 14(2), 178–192.

Tretyakov, K. (2012). Matplotlib-venn. Functions for plotting area-proportional two- and three-way Venn diagrams in matplotlib. Retrieved from: https://pypi.org/project/matplotlib-venn/.

Vachez, L., Teste, C., & Vanoosthuyse, V. (2022). DNA: RNA immunoprecipitation from S. pombe cells for qPCR and genome-wide sequencing. In R-Loops: Methods and Protocols (pp. 411–428). New York, NY: Springer US.

Wang, H., Li, C., & Liang, K. (2022). Genome-wide native R-loop profiling by R-loop cleavage under targets and tagmentation (R-Loop CUT&Tag). In R-Loops: Methods and Protocols (pp. 345–357). New York, NY: Springer US.

Werner, M., Trauner, M., Schauer, T., Ummethum, H., Márquez-Gómez, E., Lalonde, M., … & Hamperl, S. (2025). Transcription-replication conflicts drive R-loop-dependent nucleosome eviction and require DOT1L activity for transcription recovery. Nucleic Acids Research, 53(4), gkaf109.

Wulfridge, P., Yan, Q., & Sarma, K. (2022). Targeted nuclease approaches for mapping native R-loops. In R-Loops: Methods and Protocols (pp. 373–380). New York, NY: Springer US.

Xu, W., Li, K., Li, Q., Li, S., Zhou, J., & Sun, Q. (2022). Quantitative, convenient, and efficient genome-wide R-loop profiling by ssDRIP-seq in multiple organisms. In R-Loops: Methods and Protocols (pp. 445–464). New York, NY: Springer US.

Yadav, C., Yadav, R., Nanda, S., Ranga, S., & Ahuja, P. (2024). The hidden architects of the genome: a comprehensive review of R-loops. Molecular Biology Reports, 51(1), 1095.

Zhang, X., Hao, Y., & Fu, X. D. (2022). Mapping R-loops using catalytically inactive RNaseH1 (R-ChIP). In R-Loops: Methods and Protocols (pp. 359–372). New York, NY: Springer US.

Zhang, Y., Liu, T., Meyer, C. A., Eeckhoute, J., Johnson, D. S., Bernstein, B. E., … & Liu, X. S. (2008). Model-based analysis of ChIP-Seq (MACS). Genome biology, 9, 1–9.

