## Supplementary Tables and Methods for "Cellular type is a major determinant of R-loop genomic distribution"

**Supplementary information**

The Supplementary information contains:

1. Supplementary tables;
2. Supplementary methods.
1. **Supplementary tables**

| **Sample** | **Peaks number** | **Consensus peaks number** |
| --- | --- | --- |
| **hPSCs Replicate 1 (R1)** | 5718 | 4482 |
| **hPSCs Replicate 2 (R2)** | 5246 |  |
| **HAP1 WT R1** | 6747 | 4458 |
| **HAP1 WT R2** | 7893 |  |
| **HAP1 KO HNRNPA2B1 R1** | 1397 | 950 |
| **HAP1 KO HNRNPA2B1 R2** | 1327 |  |
| **HAP1 KO YTHDF2 R1** | 1815 | 1496 |
| **HAP1 KO YTHDF2 R2** | 2150 |  |

| **Sample** | **Raw reads number** | **Alignment quality, %** | **Uniquely mapped reads number** |
| --- | --- | --- | --- |
| **HAP1 WT R1** | 11,541,673 | 98.97 | 9,223,407 |
| **HAP1 WT R2** | 10,471,917 | 98.98 | 9,218,553 |
| **HAP1 KO HNRNPA2B1 R1** | 11,430,695 | 99.16 | 11,148,879 |
| **HAP1 KO HNRNPA2B1 R2** | 10,804,960 | 99.05 | 10,526,590 |
| **HAP1 KO YTHDF2 R1** | 10,546,946 | 98.25 | 10,151,324 |
| **HAP1 KO YTHDF2 R2** | 11,722,283 | 98.33 | 11,288,485 |
| **HAP1 WT Control** | 14,618,807 | 99.74 | 14,190,485 |
| **HAP1 KO HNRNPA2B1 Control** | 20,783,352 | 99.72 | 19,629,550 |
| **HAP1 KO YTHDF2 Control** | 19,858,288 | 99.69 | 18,694,330 |

Table S1. Peak numbers for the DRIP experiments.

Table S2. Mapping statistics the DRIP samples.

2. **Supplementary methods**

***Cell lines and cell culture***

REBL-PAT hiPSCs were grown in Essential 8 medium with supplement (no. A1517001) on Matrigel-coated tissue culture flasks at 37 °C with 5% CO_2_ and passaged every 3–4 d using TrypLE Select Enzyme (no. 12563029). *HNRNPA2B1* KO HAP1 cells (Horizon Discovery, no. HZGHC007378c010), *YTHDF2* KO HAP1 cells (Horizon Discovery, no. HZGHC006678c001) and their isogenic wild type parental HAP1 cells (Horizon Discovery, no. C631) were maintained on DMEM/F12 (Gibco Life Technologies, no. 11320033) supplemented with 20% heat-inactivated fetal bovine serum containing 1% pen/strep at 37 °C in with 5% CO2. Culture medium was changed daily and the cells were passaged using trypsin every 2-3d.

***DNA-RNA Immuno-Precipitation (DRIP)***

DRIP was performed according to the already published procedure (Abakir et al, 2020). Genomic DNA was isolated from the cells using the standard phenol-chloroform protocol and dissolved in nuclease-free water or TE buffer (10 mM Tris-HCl, pH 7.5; 1 mM EDTA). The samples were sonicated in a precooled (4 °C) and degassed sonicator to obtain 300–600 bp DNA fragments. We used the following parameters for sonication using Sonics Vibra Cell CV18: 25% amplitude, sonicate with pulses of 40s on/45s off for a total of 10-15 cycles. The size of genomic DNA fragments was checked using bioanalyzer and gel electrophoresis to confirm that the fragment sizes fall into the range of 300–600 bp. For each DRIP experiment, 7-10 μg of genomic DNA was diluted in 450 μl of or TE buffer (10 mM Tris-HCl, pH 7.5; 1 mM EDTA), and 50 μl of 10× IP buffer (100 mM Na-phosphate pH 7.0 (monodibasic), 1.4 M NaCl, 0.5% Triton X-100) was added to each sample, followed by thorough mixing by pipetting up and down 5 times. Then, 4-5 μl of S9.6. Antibody (Anti-DNA-RNA Hybrid Clon S 9.6 MABE1095, Merck) were added to the IP samples. The tubes were covered and sealed with Parafilm and incubated overnight at 4 °C on a tube rotator running at a speed of 20 rounds per minute. Next day, 20 μl of Dynabeads (Invitrogen by ThermoFisher 11201D Dynabeads N-280 Sheep anti-Mouse Ig) were prepared for each sample by rinsing them 2 times with 1× IP buffer. After collecting using magnetic rack, the beads were resuspended in 20 μl of 1× IP buffer and the mixture was transferred to the tubes containing the IP samples. The samples were incubated on a running tube rotator for 2h at 4 °C. After the incubation, the samples were placed in a magnetic rack, and the liquid was carefully discarded using a micropipette followed by 2 rinses in 1× IP buffer. Next, the tubes were placed on a magnetic rack for 5 min, the liquid discarded, and 250 μl of Proteinase K buffer containing 3 μl of 25 mg/ml Proteinase K added to the beads. The tubes were sealed with Parafilm and the samples incubated on a shaker running with a speed of 200 rounds per minute at 55 °C for at least 5 h (or overnight). After the incubation, the samples briefly centrifuged and the S9.6 antibody bound DNA was isolated using PCR purification kit according to the manufacturer’s instructions.

***Library preparation and high-throughput sequencing***

The *HNRNPA2B1* KO HAP1 cells, *YTHDF2* KO HAP1 cells and WT HAP1 cells DRIP sequencing libraries were prepared according to the NEBNext Ultra II DNA Library Prep Kit for Illumina (NEB, E7645) according to the manufacturer’s guidelines. The adapters were ligated according to the manufacturer’s protocol. Adapter ligated DNA was digested with USER enzyme (a mixture of Uracil DNA glycosylase (UDG) and the DNA glycosylase-lyase Endonuclease VIII) as stated in the manufacturer’s protocol. The DNA was amplified for 12 cycles and libraries were quantified using the Qubit fluorometer (Thermo Fisher Scientific). Sequencing was performed using Illumina HiSeq 1500 generating 10 million (M) 44, 50, 60-bp single-end reads per sample.

***Bioinformatics analysis of S9.6 DRIP-seq data***

Human HAP1 datasets (WT, KO HNRNPA2B1, KO YTHDF2) (PRJNA1250978 (https://www.ncbi.nlm.nih.gov/bioproject/PRJNA1250978)) and previously published human hPSCs dataset (PRJNA474076; https://www.ebi.ac.uk/ena/browser/view/PRJNA474076?show=reads) were analysed in parallel. FastQC (Andrews et al. 2010) was performed to check the quality of each dataset. According to the obtained reports, the samples containing adapter and/or primer contamination were trimmed by Trim Galore (Krueger 2015). For the reads alignment, Bowtie2 (Langmead and Salzberg 2012) and human reference assembly 110 from Ensembl were utilised. The information on the alignment quality rate and reads numbers for each sample is shown in Supplementary Table S1. The aligned reads were filtered with samtools (Li et al. 2009). Peak calling was performed using MACS2 peak caller (Zhang et al. 2008). The detection of broad peaks was performed with the parameters: --format BAM -g hs --keep-dup all. The identification of consensus peaks was done using bedtools intersect (Quinlan and Hall, 2010). The numbers of replicate and consensus peaks are shown in Supplementary Table S2. The Venn diagrams illustrating peaks overlaps were plotted using Python package ‘matplotlib-venn’ (Tretyakov, 2012). Specific loci for all the samples were visualised using Integrative Genomics Viewer (IGV) (Thorvaldsdottir et al. 2013). The code with additional details is provided in the GitHub online repository (https://github.com/katerinaoleynikova/human_samples_paper_25).
